# The structurally conserved TELR region on shelterin protein TPP1 is essential for telomerase processivity but not recruitment

**DOI:** 10.1101/2020.12.07.412684

**Authors:** Ranjodh Sandhu, Madhav Sharma, Derek Wei, Lifeng Xu

## Abstract

In addition to mediating telomerase recruitment, shelterin protein TPP1 also stimulates telomerase processivity. Assessing the *in vivo* significance of the latter role of TPP1 has been difficult, as TPP1 mutations that perturb telomerase function tend to abolish both telomerase recruitment and processivity. We sought to separate the two activities of TPP1 in regulating telomerase by considering a structure-guided mutagenesis study on the *S. cerevisiae* telomerase-associated Est3 protein, which revealed a TELR surface region on Est3 that regulates telomerase function *via* an unknown mechanism without affecting the interaction between Est3 and telomerase (1). Here, we show that mutations within the structurally conserved TELR region on TPP1 impaired telomerase processivity while leaving telomerase recruitment unperturbed, hence uncoupling the two roles of TPP1 in regulating telomerase. Telomeres in cell lines containing homozygous TELR mutations progressively shortened to a critical length that caused cellular senescence, despite the presence of abundant telomerase in these cells. Our findings not only demonstrate that telomerase processivity can be regulated by TPP1, in a process separable from its role in recruiting telomerase to telomeres, but also establish that the *in vivo* stimulation of telomerase processivity by TPP1 is critical for telomere length homeostasis and long-term cell viability.

**Significance:** Telomerase directs the synthesis of new telomeric repeats at chromosome ends, enabling cells to overcome the end replication problem and continue to divide. The shelterin protein TPP1 interacts with telomerase, promoting both telomerase recruitment and processivity (the addition of multiple telomeric repeats after a single substrate binding event). Here we show the identification of separation-of-function mutants of TPP1 that eliminate telomerase processivity but leave the telomerase recruitment function intact. When introduced into human cells in a homozygous manner, these mutations can induce critical telomere shortening and cellular senescence. Our observations therefore provide the first demonstration that telomerase processivity, in addition to telomerase recruitment, is a key regulatory step *in vivo* for continued human cell proliferation.

## Introduction

Human telomeric DNA consists of long duplex region of tandem TTAGGG repeats terminated at a 3’ single-stranded overhang (2-4). The reverse transcriptase telomerase extends telomeres by using a short segment of its RNA subunit as template to add new repeats to telomeric overhangs (5). In most human cells capable of continuous division, a homeostatic state of telomere length is maintained by balancing the lengthening effect of telomerase and the shortening effect of nucleolytic degradation and the end replication problem (6-9). Inhibition of telomerase disrupts this balance, causing progressive telomere shortening and ultimately cellular senescence (8, 10-12).

A key regulator of telomerase is the TPP1 subunit of shelterin, a multi-subunit protein complex that associates with telomeres (13). Within the shelterin complex, TRF1 and TRF2 bind sequence-specifically to the duplex telomeric repeats (14, 15), while the POT1/TPP1 heterodimer binds to the telomeric terminal overhangs (16, 17). TIN2 simultaneously interacts with TRF1, TRF2, and TPP1 (18-21), linking the double-stranded and single-stranded regions of telomeres and spreading POT1/TPP1 along the duplex telomeric tracts. TPP1 regulates two aspects of telomerase function. First, TPP1 is essential *in vivo* for recruiting telomerase to its site of action at telomeric termini and second, in the presence of POT1, TPP1 stimulates the *in vitro* processive addition of TTAGGG repeats by telomerase to a telomeric substrate (22). Both of these activities are mediated by a group of surface residues known as the TEL patch (<u>T</u>PP1 glutamate (<u>E</u>) and leucine (<u>L</u>)-rich patch) located within the N-terminal OB-fold domain of TPP1 (23-26), as mutations in the TEL patch disrupted telomerase recruitment and also abolished the stimulatory effects on enzyme processivity. A direct interaction with telomerase is critical for both activities of TPP1, as revealed by the repression of a charge-swap mutation in the TEL patch by a compensatory charge-swap mutation in the TEN domain of the human telomerase catalytic subunit (TERT) while either mutation on its own impaired telomerase recruitment and processivity (27). TPP1-regulated telomerase function is essential for the continued proliferation of human cells, since homozygous TEL patch mutations in human iPS cells caused progressive telomere shortening and ultimately cellular senescence (28).

However, left unresolved by the above analysis was whether the inability to maintain telomeres in response to a homozygous defect in the TEL patch of TPP1 is due to a recruitment defect or a processivity defect, or both. We sought to address this by asking if there was an additional surface on TPP1 that regulated only one of these two telomerase functions. To do so, we turned our attention to the *S. cerevisiae* Est3 protein, which interacts transiently with yeast telomerase late in the cell cycle (29). It adopts a protein fold that is strikingly similar to the N-terminal OB-fold domain of TPP1 (**Fig. 1A**) even though their primary sequences are considerably different (32). A structure-guided mutagenesis of the entire Est3 surface identified two clusters of residues that are each essential for telomerase function *in vivo* (1). One cluster largely overlaps with the TEL patch on TPP1 and mediates the interaction between Est3 and telomerase, arguing for a striking level of functional and structural conservation between the Est3 and TPP1 proteins. In addition, this mutagenesis identified a second cluster of residues on the surface of Est3, named the TELR region, that also regulates telomerase function through a separate mechanism that was not determined (1).

**Figure 1.**
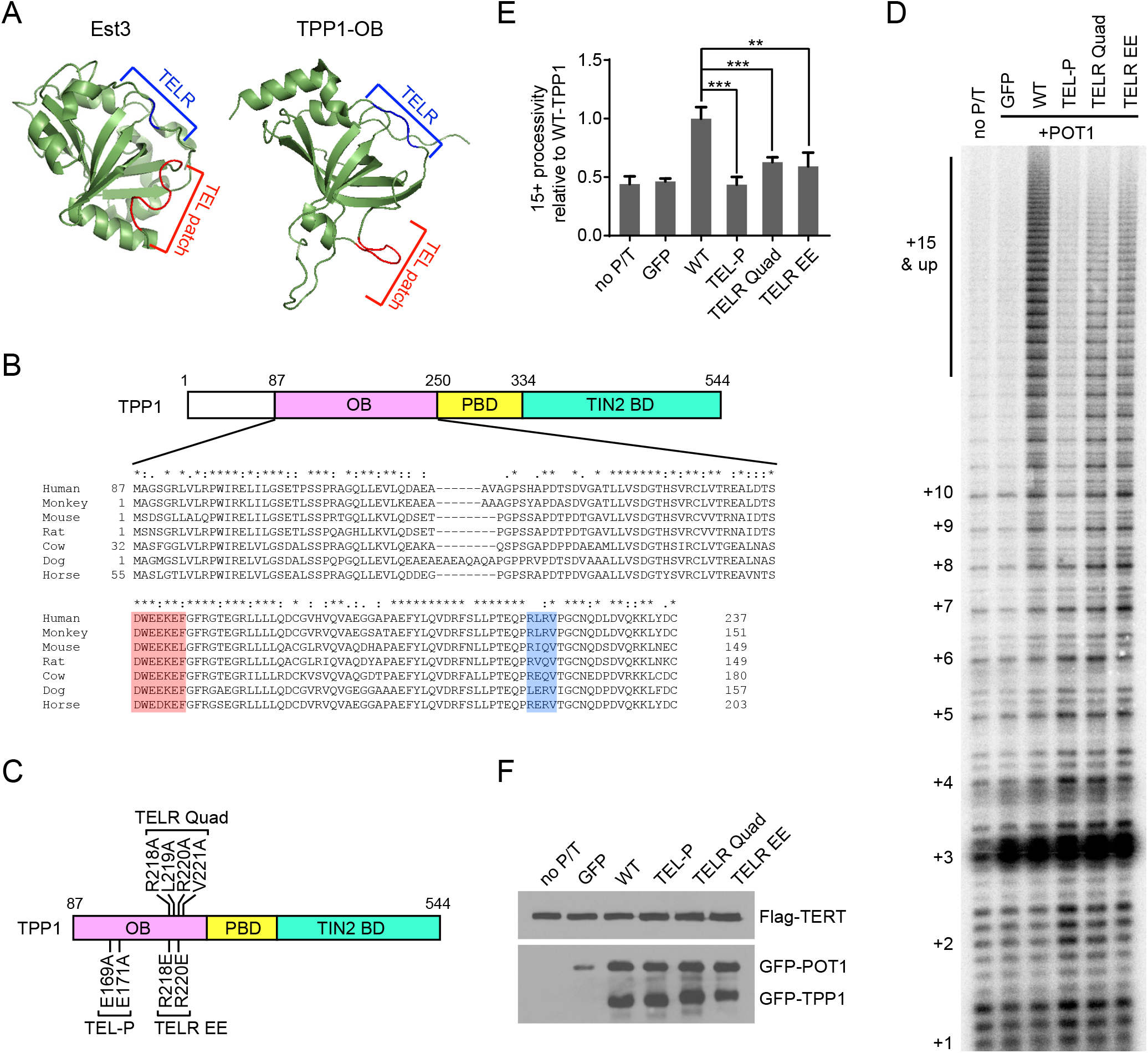
Mutations in the TELR region impaired TPP1 stimulation of telomerase processivity. (A) The TEL patch and TELR regions within Est3 and TPP1 OB-fold domain. (B) Sequence alignment of the indicated mammalian TPP1 OB-fold domains. The TEL patch acidic loop is highlighted in red and the TELR basic loop in blue. (C) Schematic of the TPP1 mutant constructs used in this study. (D) Direct telomerase activity assay using extracts from 293T cells co-transfected with expression constructs for telomerase RNA, Flag-TERT, GFP-POT1, and the indicated GFP-TPP1 alleles. “no P/T” denotes transfection without POT1 and TPP1. The number of telomeric repeats added to the oligonucleotide primer are marked along the left. (E) Quantification of telomerase processivity relative to that obtained with wildtype TPP1. Bars represent mean values of three independent experiments and SDs. P values (p<0.001 shown as *** and p<0.01 as **) were calculated by two-tailed Student’s t-tests. (F) Immunblots performed with extracts from parallel transfection of (D) to examine Flag-TERT, GFP-POT1 and GFP-TPP1 expression levels.

Here, we identified mutations in the structurally conserved TELR region on TPP1 that impair telomerase processivity without affecting recruitment of telomerase to chromosome termini. Human cell lines containing homozygous TELR mutations underwent progressive telomere shortening that led to cellular senescence, despite the presence of abundant telomerase in these cells. Our observations show that a second structural element of TPP1, in addition to the TEL patch, can control telomerase activity. Furthermore, these results establish that the *in vivo* stimulation of telomerase processivity by TPP1 is critical for telomere length homeostasis and long-term cell viability.

## Results

### Mutations in TPP1 TELR region impaired telomerase processivity

The TELR region maps to a loop connecting the β5-strand and the αC-helix of the TPP1 OB-fold (**Fig. 1A** and **1B**). We made two mutants in this region (**Fig. 1C**): the TELR Quad mutant in which four residues were mutated to alanines (R218A/L219A/R220A/V221A), and the TELR EE mutant in which two arginines were mutated to the oppositely charged glutamic acids (R218E/R220E). As a control, we also made a TEL-P mutant which contained the E169A/E171A double mutations in the TEL patch region. This mutant had been reported to disrupt TPP1’s interaction with telomerase and abolish all telomerase-associated functions of TPP1 (24). We ectopically overexpressed these mutants and examined their effects on shelterin complex assembly and telomerase processivity.

Neither of the two TELR mutants caused any detectable disruption of the shelterin complex: Immunostaining of cells transfected with plasmids for GFP-tagged TPP1 using an anti-GFP antibody followed by fluorescence in situ hybridization (FISH) using a telomeric repeat probe showed that the TELR mutations did not affect the telomeric localization of TPP1 (**Supplemental Fig. 1A**). Immunoprecipitation performed using extracts of cells co-transfected with plasmids for Flag-tagged TPP1 and GFP-tagged TIN2 or POT1 showed that the TELR mutations did not impair the interactions between TPP1 and its shelterin partners (**Supplemental Fig. 1B** and **1C**).

To determine the impact of the TELR mutations on telomerase processivity, we performed the direct telomerase activity assay using extracts of cells co-transfected with plasmids for the telomerase core subunits (TR and Flag-tagged TERT), together with plasmids for GFP-tagged TPP1 and POT1 (**Fig. 1D**). As anticipated, introduction of the TEL-P mutation completely abrogated the ability of TPP1 to stimulate the processive extension of an oligonucleotide substrate by telomerase. We observed that the TELR Quad and the TELR EE mutations also significantly reduced this ability of TPP1 (**Fig. 1D** and **1E**). Western blotting analysis of the transfected cells confirmed that the respective TERT, POT1 and TPP1 proteins were expressed at similar levels across all transfections (**Fig. 1F**).

### The TPP1 TELR region is essential for telomere length maintenance and continued cell proliferation

To assess the *in vivo* significance of the TPP1 TELR region, we used CRISPR/Cas9-mediated genome editing to generate knock-in HCT116 cells containing TELR R218E/R220E mutations (**Fig. 2A**). HCT116 is a human colon cancer cell line that has wild-type shelterin components, is telomerase-positive and karyotypically stable (30). To edit the TELR region, we transiently transfected HCT116 cells with plasmid constructs for the Cas9 protein and a TELR targeting guide RNA, together with a single-stranded oligonucleotide (ssODN) template carrying the TELR R218E/R220E mutations (**Supplemental Fig. 2A** and **2B**). We introduced translational silent mutations into the ssODN to prevent re-cutting of the edited sequence, and to create a KpnI restriction site to facilitate screening of candidate knock-in clones (**Fig. 2A, Supplemental Fig. 2B**). The TEL patch region was edited *via* a similar strategy (**Supplemental Fig. 2C** and **2D**) (31) to generate HCT116 cell lines containing the E169A/E171A mutations (**Fig. 2A**) as negative controls for telomerase-associated functions of TPP1. After the initial transfection, single colonies were picked, expanded, and screened by the KpnI digestion assay. Cell clones that showed KpnI sensitivity in the targeted genomic region were subjected to Sanger DNA sequencing to identify those that incorporated the desired mutations. For the TPP1 TELR R218E/R220E mutations, we isolated two independent homozygote clones (*TELR/TELR* clone#1 and #2); for the TEL patch E169A/E171A mutations, one homozygote clone (*TEL-P/TEL-P*) and two heterozygote clones (*TEL-P/WT* clone#1 and #2) were obtained (**Supplemental Fig. 3A** and **3B**). TPP1 protein levels in all edited clones were comparable to the parental HCT116 cells (**Supplemental Fig. 3C**), suggesting that these mutations did not significantly change TPP1 protein stability.

**Figure 2.**
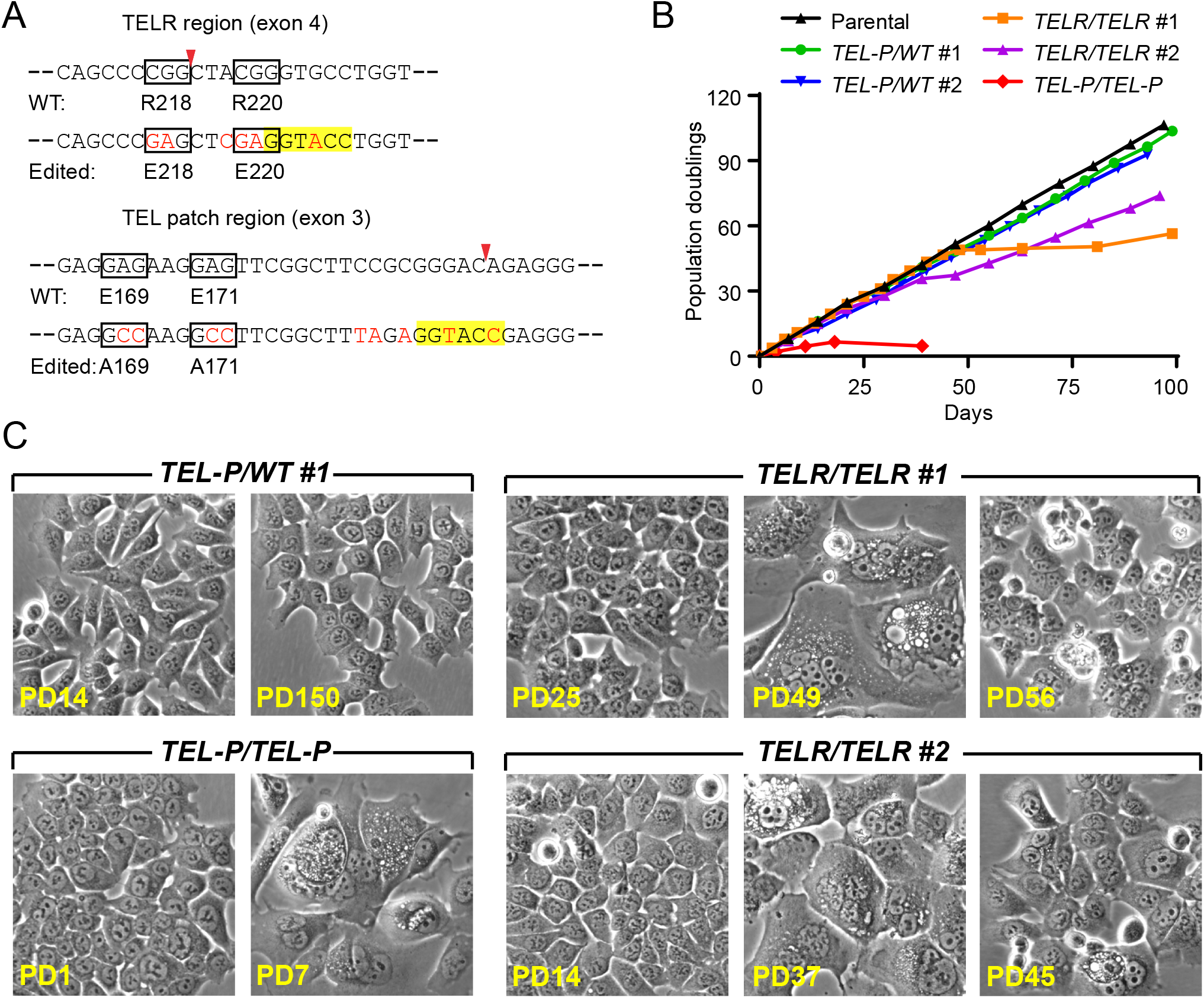
The TPP1 TELR region is essential for continued proliferation of HCT116 cells. (A) The mutagenic ssODN sequence for the TELR or TEL patch region (with the mutated nucleotides in red text) is shown below the relevant wildtype sequence of the TPP1 gene. The mutated codons were encircled with boxes and the Cas9 cut sites were marked with arrowheads. Silent mutations were introduced into the edited sequence to prevent re-cutting and to create a KpnI site (highlighted) for colony screening. (B) Growth curves of HCT116 cell lines containing homozygous TELR R218E/R220E mutations, versus those containing heterozygous or homozygous TEL patch E169A/E171A mutations. (C) Phase-contrast images of the edited cell lines at the indicated PDs. Images of pre-, peri-, or post-senescence *TELR/TELR* homozygote cells were shown.

**Figure 3.**
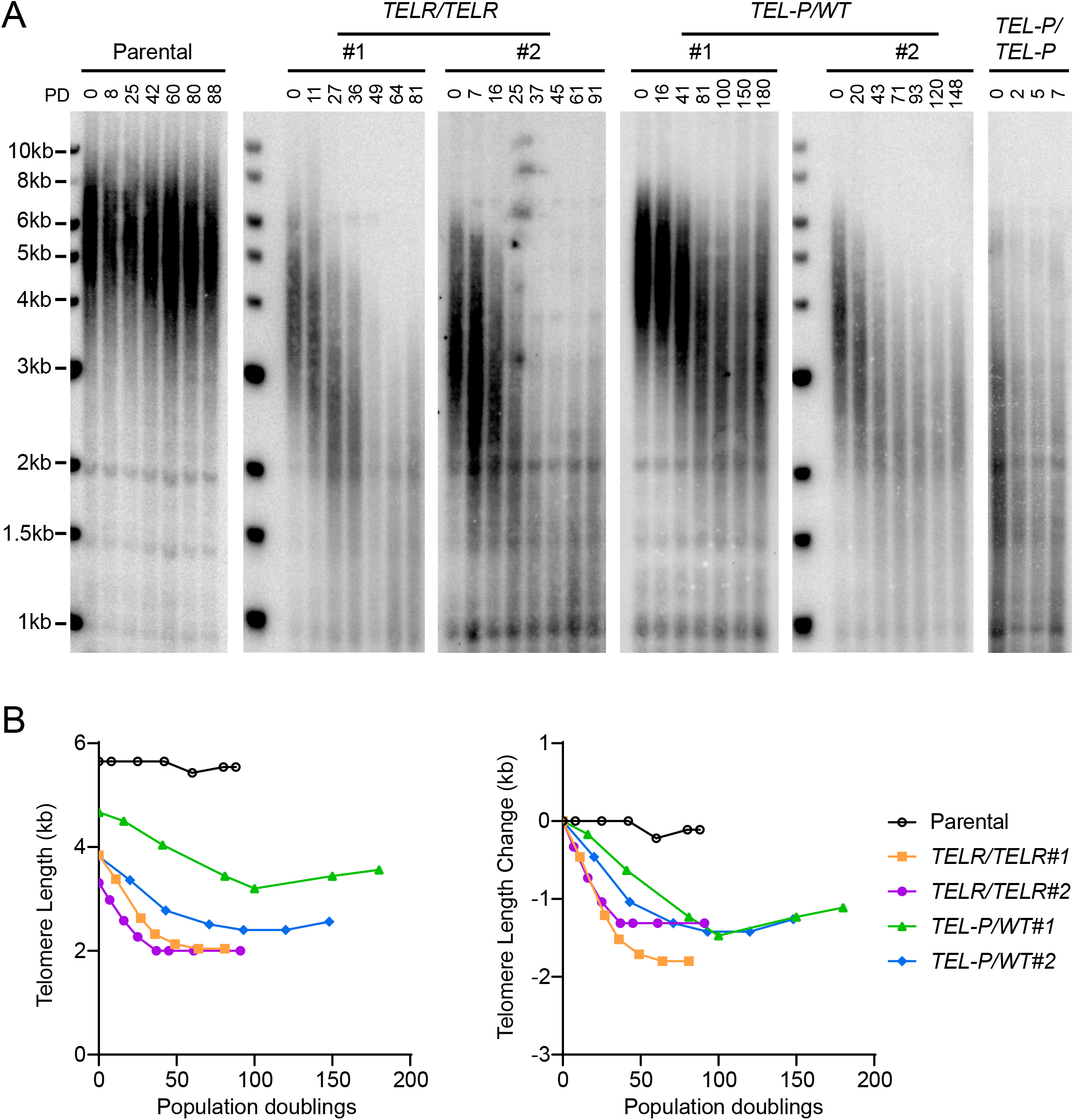
Homozygous mutations in the TPP1 TELR region caused critical telomere shortening in HCT116 cells. (A) Telomere Restriction Fragment (TRF) analysis of the edited cell lines as compared to the parental HCT116 cells over continuous passaging. (B) Left panel: Mean telomere lengths in (A) were determined by the ImageQuant software and plotted against PDs. Right panel: Quantification of changes in mean telomere length plotted against PDs.

During continuous passaging, the HCT116 parental cells proliferated at a steady rate while maintaining consistent cell morphology (**Fig. 2B and 2C**). The two clones of the *TELR/TELR* homozygote cells initially proliferated at a rate indistinguishable from the parental cells, but later entered a senescence state during which the cells became multi-nucleated, flattened, greatly enlarged and vacuolated (**Fig. 2B and 2C**). Compared to the stable telomere length in parental HCT116 cells, we observed progressive telomere shortening in the *TELR/TELR* homozygote cells (**Fig. 3A and 3B**). The onset of cellular senescence in each *TELR/TELR* clone coincided with the shortening of telomeres to a minimum mean length of ∼2kb (**Fig. 2B and 3A**), suggesting that this senescence state was induced by critical telomere shortening. Interestingly, both *TELR/TELR* homozygote clones contained a small number of cells that eventually emerged from senescence (the growth stall period was ∼30 days for clone #1 and ∼6 days for clone #2) (**Fig. 2B**). Telomere lengths remained relatively short in the post-senescence *TELR/TELR* cells (**Fig. 3A and 3B**).

For the TEL patch E169A/E171A mutations, the two clones of the *TEL-P/WT* heterozygote cells proliferated at a rate indistinguishable from the parental cells and maintained consistent cell morphology during continuous passaging (**Fig. 2B and 2C**). Telomeres in each *TEL-P/WT* heterozygote clone underwent some initial shortening before stabilizing at a new length (**Fig. 3A and 3B**). The new homeostatic state was apparently established before telomeres reached critical length since cell proliferation was not negatively impacted (**Fig. 2B**). The *TEL-P/TEL-P* homozygote cells, in contrast, went through just a few divisions before entering senescence (**Fig. 2B**). This was not surprising since their telomeres were already quite short when we obtained the clone (**Fig. 3A and 3B**). Morphology of the *TEL-P/TEL-P* senescent cells resembled that of the *TELR/TELR* clones (**Fig. 2C**) except that we didn’t observe any cell emerging from the senescence even after >100 days of culturing (data not shown).

### Mutations in the TELR region did not affect telomerase recruitment

We next examined how telomerase recruitment was impacted in the HCT116 knock-in cells. To facilitate the detection of telomerase, we transiently transfected the respective knock-in clones with plasmids encoding telomerase RNA and the catalytic subunit (TR and TERT). We then performed immunofluorescence staining against shelterin proteins TRF1 and TRF2 to detect telomeres, followed by FISH against telomerase RNA to detect telomerase. As anticipated, telomerase was recruited to ∼85% of telomeres in parental cells, while the homozygous TEL patch mutations nearly completely abrogated telomerase recruitment (**Fig. 4A and 4B**). The homozygous TELR mutations, in contrast, did not show any negative impact on telomerase recruitment: telomerase was recruited to ∼85% and ∼89% of telomeres in the respective *TELR/TELR* clones (**Fig. 4A and 4B**). This observation, in combination with the results obtained from the direct telomerase activity assay (**Fig. 1D** and **1E**), showed that mutations in the TELR region impaired telomerase processivity while leaving telomerase recruitment unperturbed.

**Figure 4.**
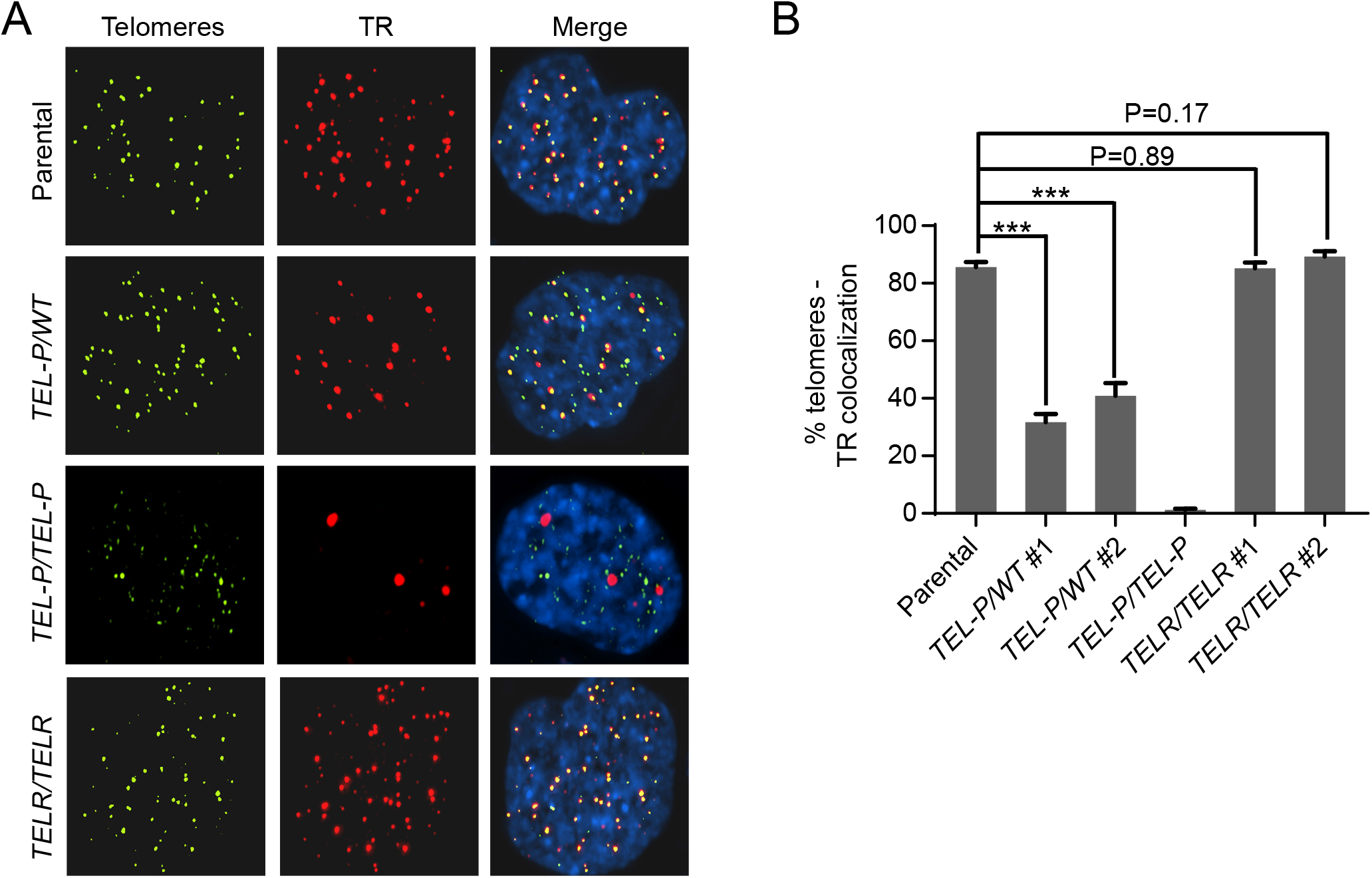
Heterozygous and homozygous TPP1 TEL patch mutations, but not the homozygous TPP1 TELR mutations, impaired telomerase recruitment in HCT116 cells. (A) Telomerase recruitment was assessed by co-localization of telomerase RNA and telomeres in each edited cell line. The cells were co-transfected with expression constructs for telomerase (TR+TERT). Immunostaining was then performed using anti-TRF1 and anti-TRF2 antibodies to detect telomeres (green), followed by RNA FISH to detect telomerase RNA (red). (B) Percentage of telomeres colocalized with telomerase RNA. Bars represent mean values of ∼50 nuclei from 10 fields of view and SEMs. All quantifications were carried out blindly. P values (p<0.001 shown as ***) were calculated by two-tailed Student’s t-tests.

We note that telomerase was only recruited to ∼32% and ∼41% of telomeres in the respective *TEL-P/WT* heterozygote clones (**Fig. 4A and 4B**). This significant reduction of telomerase recruitment caused by the heterozygous TEL patch E169A/E171A mutations provided an explanation for why these two clones reset telomere length homeostasis at a shorter length (**Fig. 3A and 3B**).

### Ectopic expression of the TPP1 TELR mutants in cultured human cells caused telomere shortening without affecting telomerase recruitment

We also examined how ectopic expressing TPP1 TELR mutants (**Fig. 1C**) affected telomere length maintenance and telomerase recruitment. We infected HCT116 cells with lentivirus expressing Flag-tagged TPP1 (**Fig. 5A**) and collected stably infected cells for bulk telomere length analysis. Overexpression of wild-type TPP1 led to robust telomere lengthening. The TELR Quad and TELR EE mutants, in contrast, each induced progressive telomere shortening (**Fig. 5B and 5C**). Similar telomere shortening was also observed in cells overexpressing the TEL patch E169A/E171A mutant (**Fig. 5B and 5C**). Unlike the *TEL-P/TEL-P* and *TELR/TELR* homozygotes, HCT116 cells overexpressing the TPP1 TEL-P and TELR mutants had their telomeres stabilized at a mean length slightly above 3kb and continued to proliferate at a constant rate (data not shown), most likely due to the presence of wild-type TPP1 expressed from its endogenous locus.

**Figure 5.**
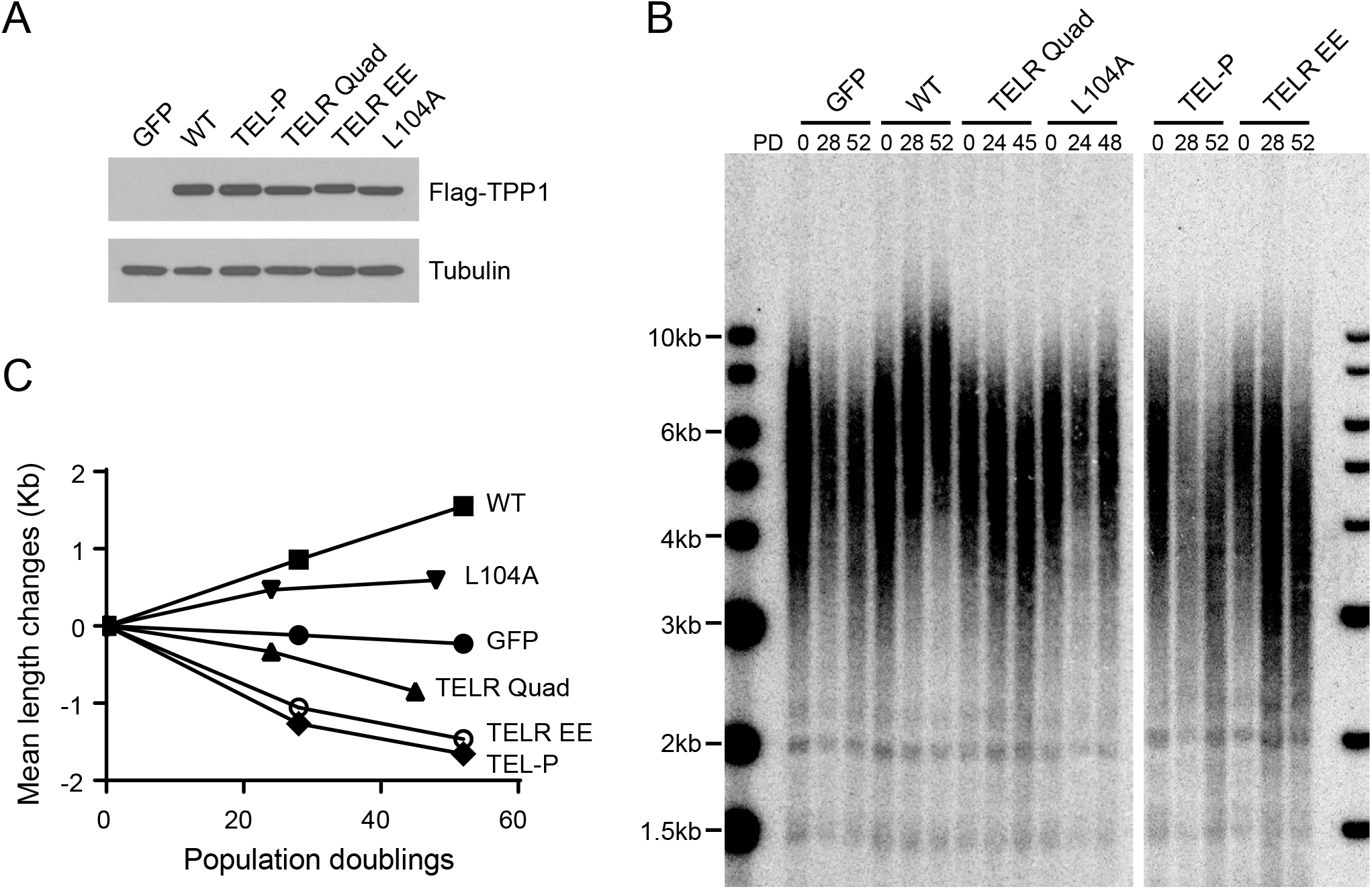
Ectopic expression of the TPP1 TELR mutants in HCT116 cells led to progressive telomere shortening. (A) Lentiviral expression level of Flag-tagged TPP1 alleles was examined by immunoblotting using an anti-Flag antibody. Tubulin was included as the loading control. (B) Telomere Restriction Fragment (TRF) analysis of cells expressing the indicated TPP1 constructs over continuous passaging. HCT116 Cells were infected with lentiviruses expressing Flag-tagged TPP1. The infected cells were pooled, continuously passaged, and collected at indicated population doublings (PDs). (C) Mean telomere length changes were plotted against population doublings.

To examine how ectopic expression of the TELR mutants affected telomerase recruitment, we transiently transfected HeLa1.2.11 cells (a HeLa subclone with long telomeres frequently used for immunofluorescence study of telomeric proteins) with plasmids encoding telomerase core subunits (TR and TERT) and GFP-tagged TPP1 variants. We then performed immunofluorescence staining against GFP-TPP1, followed by FISH against telomerase RNA. As all the TPP1 variants had been shown to localize normally to telomeres (**Supplemental Fig. 1A**), telomerase recruitment was assessed by quantifying the co-localization between telomerase RNA and the GFP-TPP1 foci. Our data showed that neither the TELR Quad nor the TELR EE mutant induced any adverse effect on telomerase recruitment: telomerase was detectable at ∼90% of the GFP-TPP1 foci in cells expressing these mutants, comparable to the level in cells expressing wild-type TPP1 (**Supplemental Fig. 4A and 4B**). Overexpression of the TEL-P mutant, as expected, sharply decreased telomerase recruitment to ∼10% of the GFP-TPP1 foci (**Supplemental Fig. 4A and 4B**). Results of the ectopic expression studies further corroborated our findings from the homozygous *TELR/TELR* mutant cell lines: the TELR region, although not required for telomerase recruitment, is essential for telomerase to extend telomeres.

We also examined the telomere length effect of a TPP1 L104A mutant. L104A was previously reported to significantly decrease TPP1 stimulation of telomerase processivity but not telomerase recruitment (24, 28, 32), similar to the TELR mutations. We found that overexpression of TPP1 L104A in HCT116 cells led to steady telomere extension, although the extension rate was less dramatic than that induced by the wild-type TPP1 (**Fig. 5B and 5C**). Our observation is in agreement with another overexpression study showing TPP1 L104A caused telomere lengthening in HeLa cells (32). We speculate that, compared to the TELR mutants, the L104A mutant may still retain a partial telomerase regulating function *in vivo*. Indeed, in human iPS cells harboring a deleterious homozygous TEL patch mutation, expression of the TPP1 L104A cDNA from an engineered AAVS1 genomic locus was found to prevent critical telomere shortening and rescue cell viability (28).

### Ectopic expression of telomerase rescued *TELR/TELR* but not *TEL-P/TEL-P* homozygotes from critical telomere shortening-triggered cellular senescence

We next investigated whether telomere maintenance and cell viability in the *TELR/TELR* and *TEL-P/TEL-P* homozygotes could be rescued by ectopic expressing wild-type TPP1 or telomerase. We infected respective pre-senescence cells with lentivirus expressing wild-type TPP1 or the core components of telomerase (TERT+TR). Stably infected cells were then pooled, continuously passaged, and collected for cell counts and bulk telomere length analysis.

Overexpression of wild-type TPP1 led to almost identical levels of telomere lengthening and the bypass of senescence in *TELR/TELR* and *TEL-P/TEL-P* cells (**Fig. 6A and 6B**), suggesting that the TELR mutant, similar to the TEL patch mutant, does not have a dominant-negative effect over wild-type TPP1’s ability in promoting telomerase function. Overexpression of telomerase, in contrast, extended telomeres in the *TELR/TELR* but not the *TEL-P/TEL-P* cells (**Fig. 6A**).

**Figure 6.**
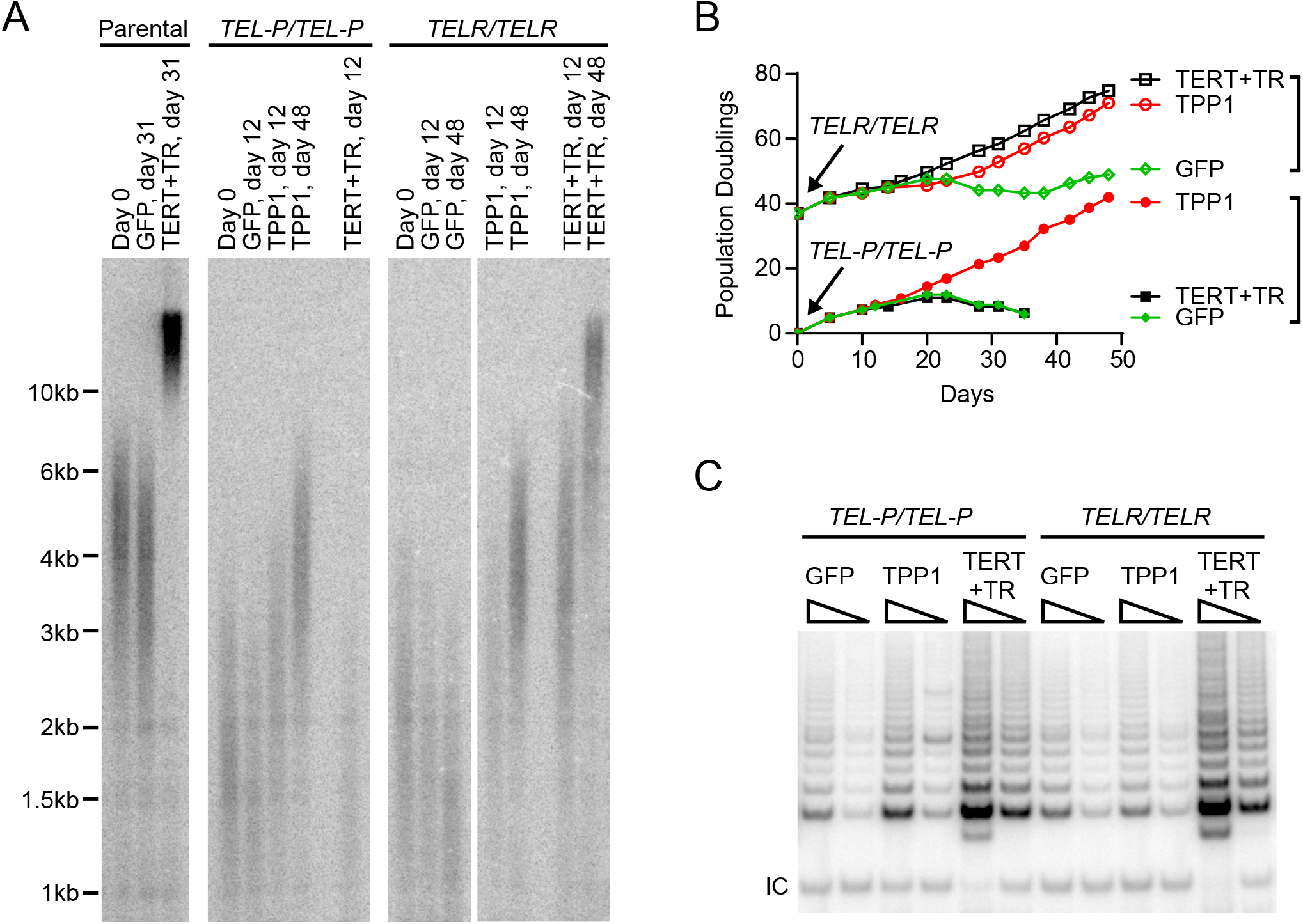
Telomerase expression prevented the *TELR/TELR* homozygotes, but not the *TEL-P/TEL-P* homozygotes, from entering the critical telomere shortening-induced cellular senescence. (A) Telomere Restriction Fragment analysis of the *TELR/TELR* or *TEL-P/TEL-P* homozygotes overexpressing GFP, TPP1, or telomerase. Pre-senescence *TELR/TELR* (at PD 37) and *TEL-P/TEL-P* cells (at PD 0) were infected with lentivirus expressing a GFP control, Flag-tagged TPP1, or telomerase (TERT+TR). Infected cells were pooled, continuously passaged, and collected for telomere length analysis and cell counts. (B) growth curves of the *TELR/TELR* or *TEL-P/TEL-P* homozygotes overexpressing GFP, TPP1 or telomerase. (C) Telomerase enzymatic activity examined by the TRAP assay. Whole cell extracts from 100 and 25 cells were analyzed for each cell line. IC: internal PCR control.

Conceivably, *TELR/TELR* but not *TEL-P/TEL-P* homozygotes were rescued from the critical telomere shortening-triggered senescence by telomerase overexpression (**Fig. 6B**). TRAP analysis showed that overexpressing telomerase increased telomerase activity to comparable levels in *TELR/TELR* and *TEL-P/TEL-P* homozygotes (**Fig. 6C**), suggesting that the failed rescue in the latter cells cannot be attributed to a lack of core telomerase enzymatic activity.

## Discussion

Although TPP1 has long been known to stimulate telomerase processivity *in vitro*, its potential *in vivo* importance remained unknown due to the inability to cleanly separate this role of TPP1 from that in recruiting telomerase. In this study, we show that the TPP1 TELR mutants significantly impaired telomerase processivity while leaving the telomerase recruitment function intact. Using CRISPR/Cas9-mediated genome editing technology, we generated HCT116 knock-in cell lines harboring homozygous TPP1 TELR mutations. Telomeres in the edited cells progressively shortened. When they reached a critical length, the cells entered a senescence state. These results therefore constitute the first demonstration that TPP1-stimulated telomerase processivity is essential for telomere length maintenance and long-term cell viability. Furthermore, our data provide direct evidence that telomerase processivity can be regulated by TPP1 in a process separable from its role in recruiting telomerase. This suggests that telomerase processivity, in addition to telomerase recruitment, maybe a key regulatory step for *in vivo* telomerase function.

One potential concern is that the TELR mutations might have conveyed a partial defect in TPP1-telomerase interaction that our telomerase recruitment assay was not sensitive enough to detect. We note that compared to the *TELR/TELR* mutant cells, the *TEL-P/WT* cells containing heterozygous mutations in the TEL patch manifested a much more significant reduction in telomerase recruitment (**Fig. 4A and 4B**) yet less severe telomere shortening phenotype (**Fig. 3A and 3B**). These observations argue that the telomere maintenance defect associated with the TELR mutations is not due to a slight decrease in telomerase recruitment, but rather a deficiency in a different aspect of TPP1’s telomerase-associated function. The exact mechanism by which the TELR region impacts telomerase processivity remains to be determined. Recent structural and single molecule studies of telomerase suggest that the TEN-RBD-RT-TRAP ring on TERT forms DNA substrate binding motifs to aid template translocation after the addition of each telomeric repeat (33-35). We note the conservation of one or two arginines within the short TELR region across mammalian TPP1 alleles (**Fig. 1B**). Whether these residues contribute to the formation of a basic DNA-binding pocket together with the TERT ring to help retain telomeric DNA substrate during template translocation will be a subject for future investigation.

In both clones of the *TELR/TELR* homozygote cells, after an extended period of stalled cell proliferation, we observed the outgrowth of a small subset of cells. Compared to the original cell population, the survivor cells exhibited increased telomerase enzymatic activity as indicated by the TRAP assay results (**Supplemental Fig. 5A and 5B**). This suggests that the bypass of the senescence was prompted by telomerase-dependent but not ALT-dependent telomere maintenance. In line with this observation, overexpression of telomerase in *TELR/TELR* homozygotes was found to extend telomeres and sustain cell proliferation (**Fig 6A and 6B**). Although the TELR mutations impaired telomerase processivity, telomerase recruitment was normal in the *TELR/TELR* cells. Overexpression of telomerase produced excess amount of telomerase molecules. The decreased length of extension per telomerase binding event could be compensated by multiple rounds of extension by different telomerase molecules. As a result, telomeres were extended in these cells. Overexpression of telomerase in the *TEL-P/TEL-P* mutant cells, in contrast, failed to extend telomeres. This is expected since telomerase recruitment was completely abrogated in the *TEL-P/TEL-P* cells.

Lastly, our data suggest that the TELR equivalent region on *S. cerevisiae*’s Est3 may likewise contribute to telomere length maintenance through enhancing telomerase processivity. Although significant differences exist in the composition and structure of telomerase accessory factors in budding yeasts and mammals, this study further highlights the possibility that telomerase action at telomeres is regulated through some conserved modes during evolution.

## Materials and Methods

### Cell lines

The HCT116 parental cells were obtained from ATCC. HeLa1.2.11 cells were kindly provided by Dr. Titia de Lange. All cell lines were grown in DMEM supplemented with 10% fetal bovine serum.

### Plasmids

The pSpCas9(BB)-2A-Puro (PX459) vector (Addgene plasmid #62988) (36) and the gRNA cloning vector (Addgene plasmid #41824) (37) were gifts from Drs. Feng Zhang (Broad Institute) and George Church (Harvard University), respectively. The pHR’CMV lentiviral expression system was kindly provided by Dr. Didier Trono (EPFL). Flag-TPP1, GFP-TPP1, GFP-POT1, and Flag-hTERT lentiviral constructs contain respective cDNA driven by a CMV promoter, followed by an internal ribosome entry site and a hygromycin resistance gene. For full-length TPP1 expression, TPP1 coding sequence from a.a. 87-544 as previously defined (24, 38) was used. Telomerase RNA (hTR) expression was driven by an IU1 promoter (39). Lentivirus was prepared as described previously (40).

### CRISPER-Cas9 mediated genome editing

Guide RNAs targeting the TPP1 genomic locus were designed using Feng Zhang Lab (Broad Institute) open access Guide Design Tool (crispr.mit.edu). For each set of mutations, three selected guide RNAs based on their high Guide Design Tool scores and low off-target probabilities were assessed by the T7 endonuclease I (NEB) assay per manufacturer’s direction. The top ranked guide RNAs were chosen for subsequent genomic editing. Their targeting sequences were as follows: (TELR targeting) CAACCAGGCACCCGTAGCCG; (TEL patch targeting) GTTCGGCTTCCGCGGGACAG. ssODNs containing ∼70bp of flanking sequence on each side of the targeted mutations were used to introduce the desired mutations into the TPP1 genomic locus. Silent mutations were included in each ssODN sequence to block re-cutting of the edited sequence and to generate a KpnI restriction site to facilitate the identification of edited cell clones.

To edit the TPP1 genomic locus, HCT116 cells were transiently transfected by a plasmid expressing Cas9 and one expressing both the puromycin-resistance gene and guide RNA, together with the ssODN using the jetPrime reagent (Polyplus transfection). One day after transfection, 2μg/ml puromycin was added and kept in culture media for about two days to select out the non-transfected cells. Three days later, cells were trypsinized and a fraction of cells were seeded at 300 cells/10cm-plate for colony formation. The remaining cells were collected, their genomic DNA extracted, and used for assessing editing efficiency *via* the KpnI digestion assay as described below. ∼2 weeks after seeding, individual colonies were picked, expanded, and screened by the KpnI digestion assay followed by Sanger DNA sequencing to identify those clones that incorporated the desired mutations.

### Screening of knock-in clones

We adopted a KpnI digestion screen strategy (31) to identify cell clones that had incorporated a KpnI restriction site at the targeted TPP1 genomic locus *via* ssODN-mediated homology-directed repair. Briefly, genomic DNA was extracted from each single colony of cells, and then the respective target genomic mutagenesis region was PCR amplified and subjected to KpnI restriction digestion. PCR primers and the anticipated results from the edited clones are as follows: (TELR region) Forward-GGACCCACAGTGTCCGATG, Reverse-TACCTGCATTGGACGAGGTG, PCR product size ∼730bp, digested product sizes ∼310+420bp; (TEL patch region) Forward-GCTTGAGGTGAGCCCCCATT, Reverse-CCTACCCCATAGGCGTCTGC, PCR product size ∼670bp, digested product sizes ∼360bp+310bp. Cell clones that showed KpnI sensitivity were then expanded. To identify those clones that incorporated the desired mutations, respective target genomic mutagenesis region on TPP1 was PCR amplified and subjected to Sanger DNA sequencing using the following primers: Forward-GCTTGAGGTGAGCCCCCATT, Reverse-GACCATGGGATGAGTCAAGGCTT. The PCR product is ∼1030bp and spans both the TELR and the TEL patch region.

### Direct telomerase activity assay

Cell extracts were prepared from 293T cells transiently transfected with hTERT, hTR, TPP1 and POT1 expressing plasmids as previously described (24). Direct telomerase activity assay was carried out according to an established protocol (31, 41). Briefly, each 20 μl reaction contained 50 mM Tris-Cl (pH8.0), 30 mM KCl, 1 mM MgCl2, 1 mM spermidine, 5 mM β-mercaptoethanol, 1 μM primer a5 (TTAGGGTTAGCGTTAGGG), 500 μM dATP, 500 μM dTTP, 2.92 μM unlabeled dGTP, 0.33 μM radiolabeled dGTP (3000 Ci/mmol), and 3 μL of cell extracts. Reactions were incubated at 30 °C for 45 min. DNA was precipitated and resolved on 10% acrylamide, 7 M urea, and 1xTBE sequencing gels. Gels were scanned with a Phosphorimager and analyzed by the ImageQuant software (GE Healthcare). Processivity was calculated using the “15+ method” as described previously (24).

### Antibodies

Immunoprecipitation and immunoblotting were carried out according to standard protocols with the following antibodies: rat anti-Flag L5 antibody (BioLegend; 637301), rabbit anti-GFP antibody (Novus; NB600-308), mouse anti-TPP1 (Abnova; H00065057-M02). Immunofluorescence staining and FISH were carried out as described previously (42) with the following antibodies: rabbit anti-GFP antibody (Novus; NB600-308), mouse anti-TRF1 antibody (GeneTex; GTX70304). mouse anti-TRF2 antibody (Millipore; 05-521), rabbit anti-coilin antibody (GeneTex; GTX112570).

### Telomere restriction fragment analysis

4μg of each genomic DNA was digested with HinfI and RsaI and separated on 0.7% agarose-TBE gels. After depurination and denaturation, DNAs were transferred to a Hybond XL membrane and hybridized to a radiolabeled telomeric probe (CCCTAA)_4_. Blots were imaged on a PhosphorImager and analyzed by the ImageQuant software (GE Healthcare). Mean telomere lengths were calculated according to the positions of radiolabeled 1kb DNA ladder (NEB) run on the same gel.

### Telomerase recruitment assay

Cells were transiently transfected with plasmid constructs for hTERT and hTR using Lipofectamine 2000 (Life Technologies) or the jetPrime reagent (Polyplus transfection). 24 hours later, transfected cells were trypsinized and seeded onto sterile coverslips placed in 6-well plate. Two days after seeding, combined immunofluorescence staining-telomerase RNA FISH was carried out as described (42). For immunostaining of telomeres, the cells were incubated with anti-TRF1 and anti-TRF2 primary antibodies, followed by an Alexa Fluor 488 (Molecular Probes) conjugated secondary antibody. For immunostaining of GFP-TPP1, an anti-GFP primary antibody was used. Subsequent telomerase RNA FISH was performed with a mixture of three Cy3-conjugated telomerase RNA probes (43). Cell images were acquired using a Nikon Ti-U microscope with a 100x objective and collected as a stack of 0.2 μm increments in the z-axis. After deconvolution using the AutoQuant X3 software (Media Cybernetics), images were viewed with the Maximal Projection option on the z-axis. All image files were randomly assigned coded names to allow blinded scoring for spots co-localization.

### TRAP telomerase activity assay

Telomerase enzymatic activity was analyzed using the TRAPeze kit (Millipore) per manufacturer’s directions. The telomeric extension products were separated by 10% TBE-PAGE and visualized by a PhosphorImager (GE Healthcare). TRAP products intensity in each lane were quantified by the ImageQuant Software and normalized to the respective internal control intensity.

## Acknowledgments

We thank Dr. Joshua Meckler for performing preliminary experiments on guide RNA assessment; Dr. Deborah Wuttke for help with X-ray structure interpretation; Drs. Deborah Wuttke and Vicki Lundblad for critical reading of the manuscript. This study was supported by grants from the American Cancer Society [RSG-12-069-01-DMC] and the University of California Cancer Research Coordinating Committee (CRCC) to L.X.

## Author Contributions

R.S. and L.X. conceived the experiments. R.S., M.S., and D.W. performed the experiments and analyzed the data. R.S. and L.X. wrote the manuscript.

---

**Supplementary Figure 1.**
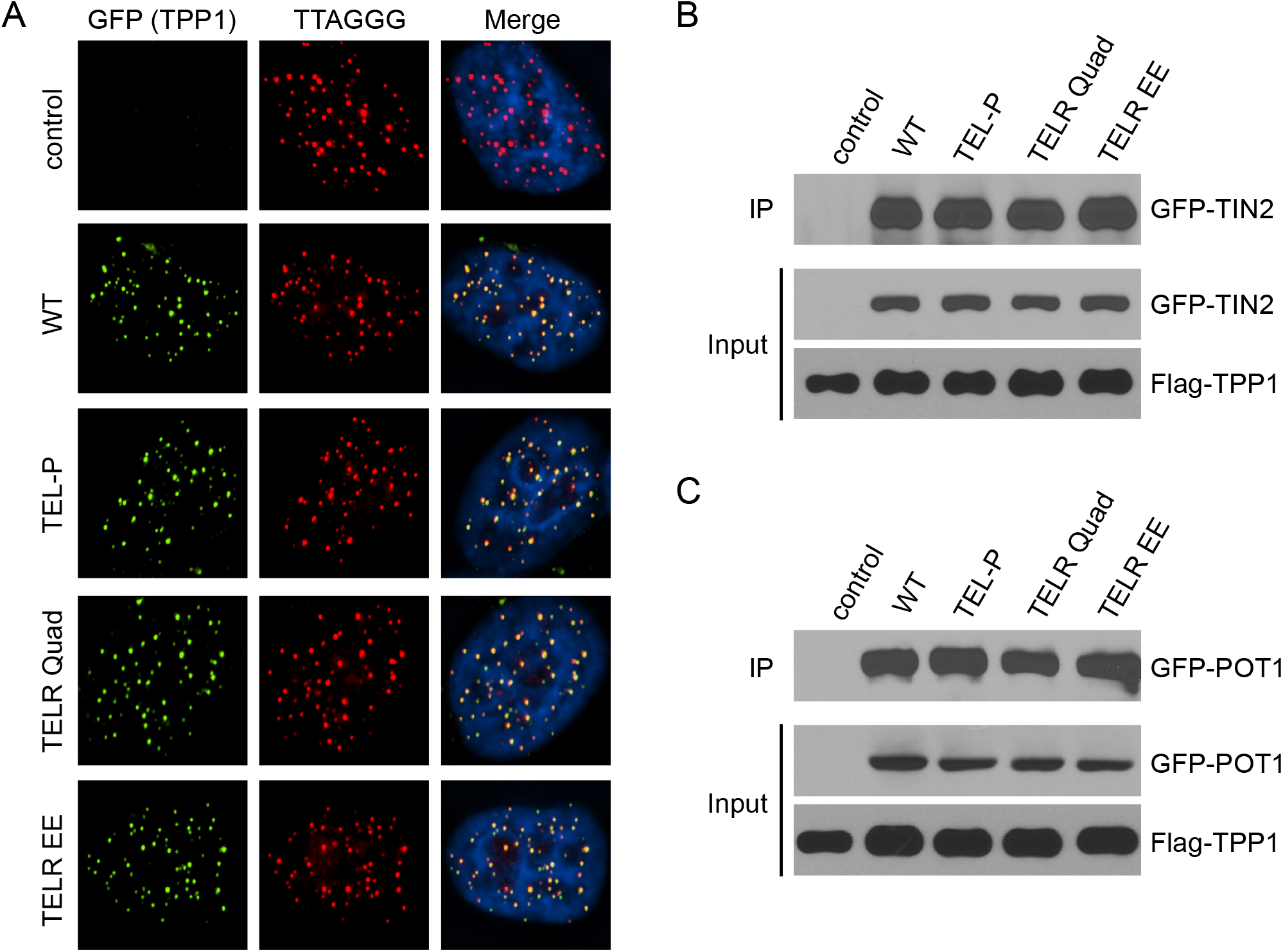
Mutations in the TELR region of TPP1 do not affect its localization to telomeres nor its interaction with other shelterin proteins. (A) Telomeric localization of GFP-tagged TPP1 examined by immunostaining using an anti-GFP antibody followed by FISH using a telomeric probe. (B) The interaction between TPP1 and its shelterin binding partner TIN2 or (C) POT1 examined by co-immunoprecipitation analysis. Cells were transfected with plasmids encoding the indicated Flag-tagged TPP1 variants together with GFP-tagged TIN2 or GFP-tagged POT1. Co-immunoprecipitation of the whole cell extracts was performed with an anti-Flag antibody, followed by immunoblotting with an anti-GFP antibody.

**Supplementary Figure 2.**
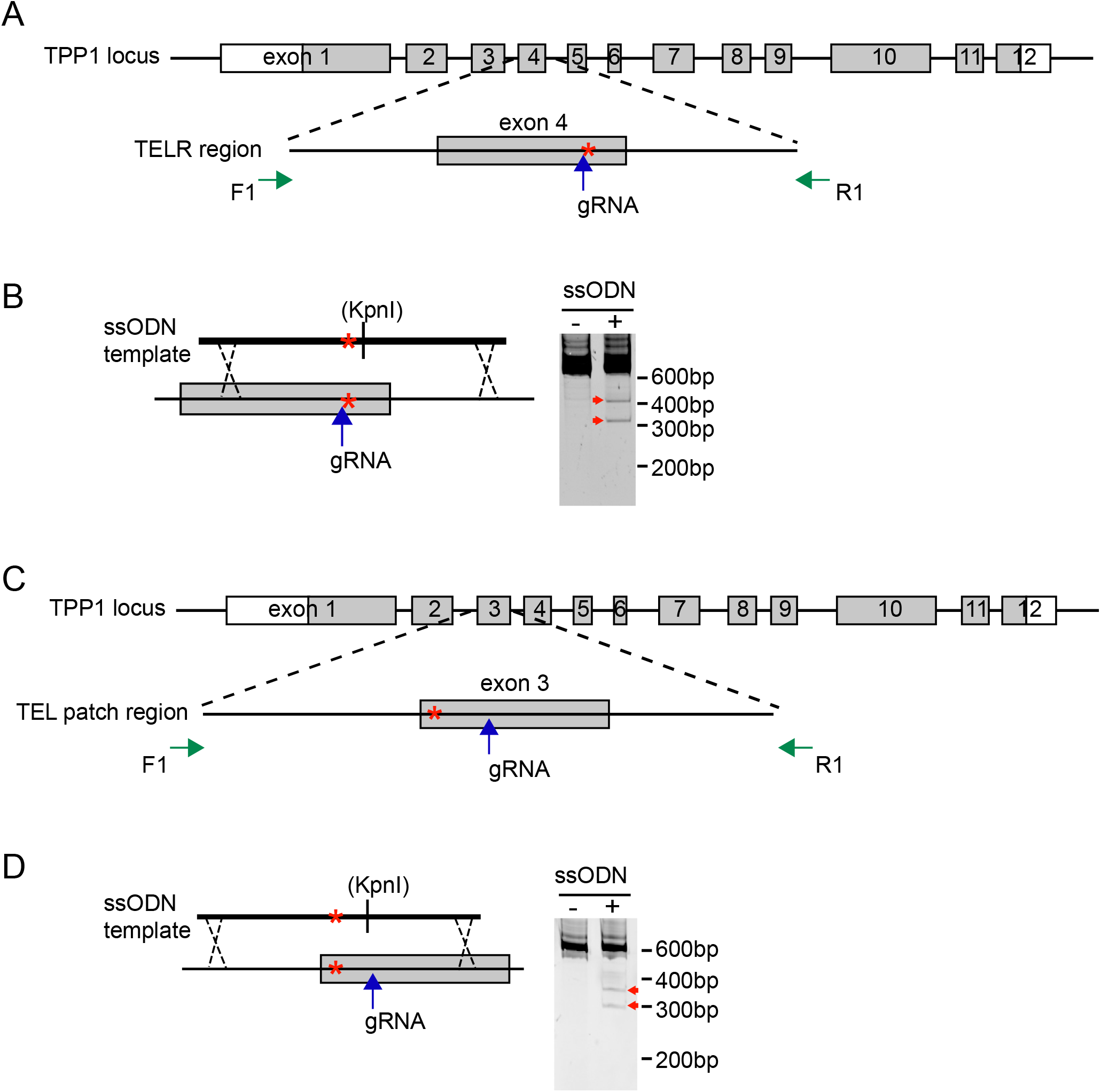
Targeting strategy for knock-in mutations in the TELR or the TEL patch region of TPP1 *via* CRISPR/Cas9-mediated genome editing. (A) Schematic of the TELR region guide RNA targeting site (marked by a red asterisk). Exons are represented as boxes and introns as lines. Coding region of exons are shaded in gray. (B) (Left) Schematic of the single-stranded oligonucleotide (ssODN) template for knocking in mutations in the TELR region. Silent mutations were included in the ssODN to prevent re-cutting of the edited sequence and to generate a KpnI site for colony screening. (Right) Knock-in efficiency evaluated by KpnI digestion of the PCR products (primers marked by green arrows) amplified from bulk genomic DNA of cells transiently transfected with plasmids for Cas9 and guide RNA, with or without the ssODN. Red arrows indicate the predicted KpnI cleavage products. (C) Schematic of the TEL patch region guide RNA targeting site (marked by a red asterisk). (D) (Left) Schematic of the single-stranded oligonucleotide (ssODN) template for knocking in the TEL patch E169A/E171A mutations. Silent mutations were included in the ssODN to prevent re-cutting of the edited sequence and to generate a KpnI site for colony screening. (Right) Knock-in efficiency evaluated by KpnI digestion of the PCR products (primers marked by green arrows) amplified from bulk genomic DNA of cells transiently transfected with plasmids for Cas9 and guide RNA, with or without the ssODN. Red arrows indicate the predicted KpnI cleavage products.

**Supplementary Figure 3.**
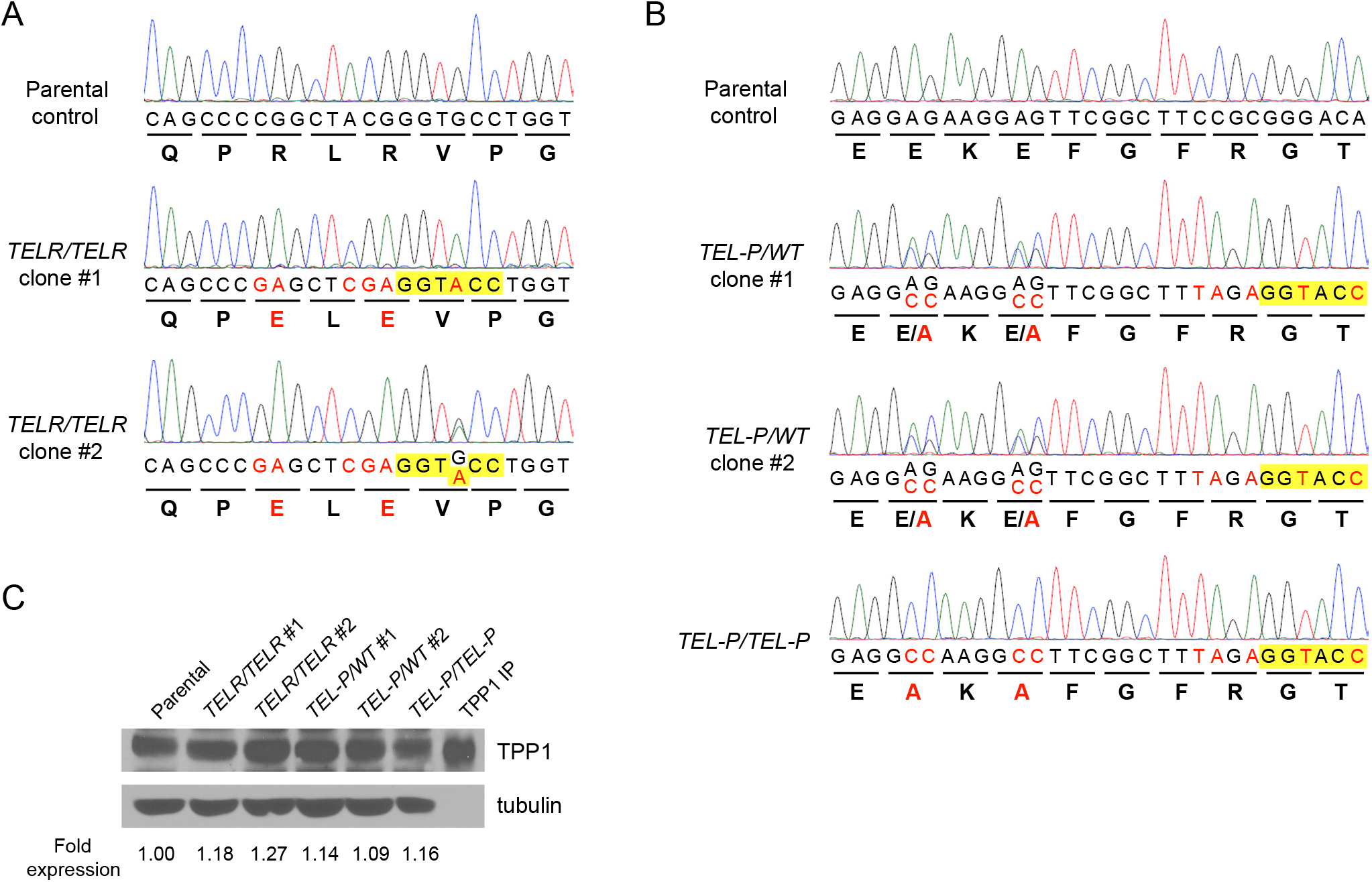
Validation of the HCT116 knock-in cell lines containing mutations in the TELR or TEL patch region. (A) Sequence verification of the TELR homozygote cell lines (*TELR/TELR* clone #1 and #2) containing the R218E(CGG→GAG)/R220E(CGG→GAG) mutations. Note: clone #2 is homozygous for the TELR mutations but heterozygous for the KpnI site (B) Sequence verification of the TEL patch heterozygote (*TEL-P/WT*#1 and #2) and homozygote mutant lines (*TEL-P/TEL-P*) containing the E169A(GAG→GCC)/E171A(GAG→GCC) mutations. (C) TPP1 expression levels in the indicated HCT116 knock-in cell lines examined by immunoblotting, with tubulin serving as a loading control. Pull down of TPP1 from parental cells by an anti-TIN2 antibody (the lane marked as TPP1 IP) was included as a positive control for the TPP1 band. Fold of TPP1 expression was quantified by the ImageJ software and normalized to tubulin levels.

**Supplementary Figure 4.**
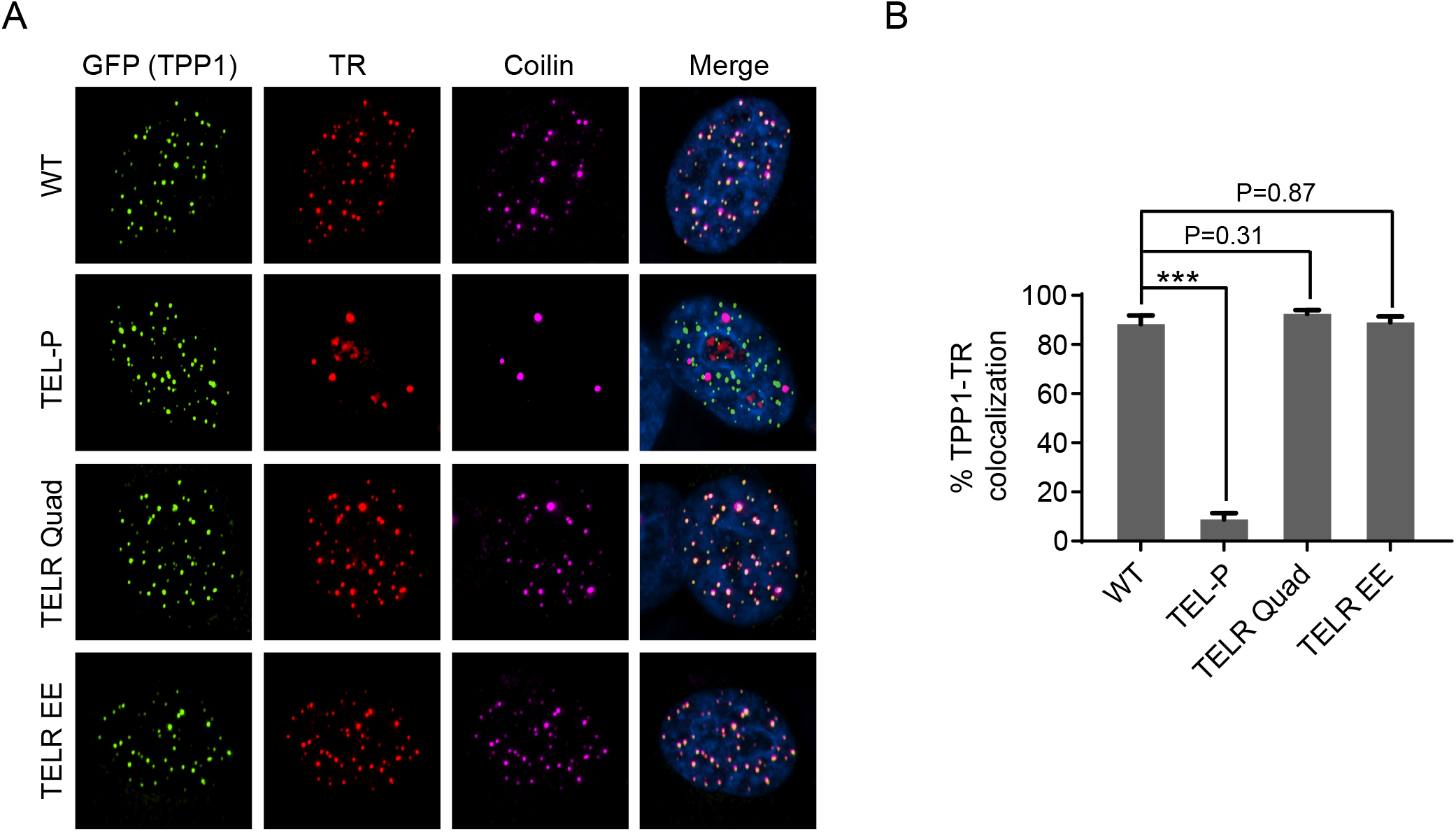
Overexpression of the TPP1 TEL patch mutant, but not any of the TELR mutants, impaired telomerase recruitment to telomeres. (A) Telomerase recruitment to telomeres in HeLa1.2.11 cells overexpressing TPP1 variants examined by co-localization of telomerase RNA and telomeres. HeLa1.2.11 cells were co-transfected with expression constructs for telomerase RNA, Flag-TERT, and the indicated GFP-TPP1 variants. Immunostaining was performed with an anti-GFP antibody to detect GFP-TPP1 (green), and an anti-coilin antibody to detect Cajal bodies (purple), followed by RNA FISH to detect telomerase RNA (red). (B) Quantification of telomerase RNA and GFP-TPP1 colocalization. Bars represent mean values of ∼50 nuclei from 10 fields of view and SEMs. All quantifications were carried out blindly. P values (p<0.001 shown as *** and p<0.01 as **) were calculated by two-tailed Student’s t-tests.

**Supplementary Figure 5.**
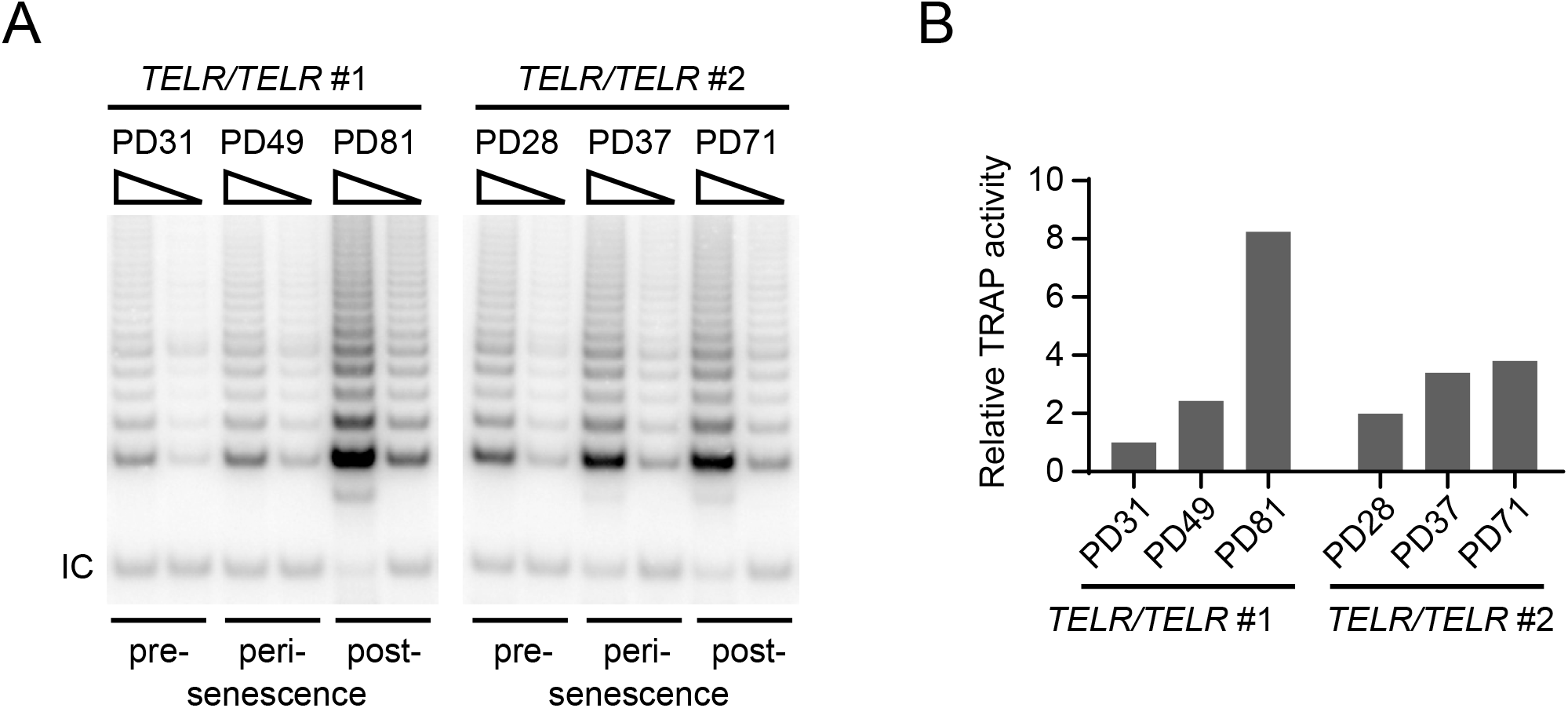
Post-senescence *TELR/TELR* survivor cells had increased telomerase enzymatic activity. (A) *in vitro* telomerase activity in pre-, peri-, or post-senescence *TELR/TELR* homozygote cells examined by TRAP assay. Whole cell extracts from 100 and 25 cells were analyzed for each cell line. IC: internal PCR control. (B) Telomerase TRAP activity in the indicated cells normalized to that in pre-senescence clone #1 cells. Quantification was conducted using readout from 25 cells for each line.

